# High-throughput single-cell, single-mitochondrial DNA assay using hydrogel droplet microfluidics

**DOI:** 10.1101/2024.01.29.577854

**Authors:** Juhwan Park, Parnika S. Kadam, Yasemin Atiyas, Bonirath Chhay, Andrew Tsourkas, James H. Eberwine, David A. Issadore

**Affiliations:** Department of Bioengineering, School of Engineering and Applied Science, University of Pennsylvania, Philadelphia, Pennsylvania 19104, USA; Department of Pharmacology, Perelman School of Medicine, University of Pennsylvania, Philadelphia, Pennsylvania 19104, USA

**Keywords:** Droplet microfluidics, High-throughput, Rolling circle amplification, Single-cell, Single mitochondrial DNA

## Abstract

There is growing interest in understanding the biological implications of single cell heterogeneity and intracellular heteroplasmy of mtDNA, but current methodologies for single-cell mtDNA analysis limit the scale of analysis to small cell populations. Although droplet microfluidics have increased the throughput of single-cell genomic, RNA, and protein analysis, their application to sub-cellular organelle analysis has remained a largely unsolved challenge. Here, we introduce an agarose-based droplet microfluidic approach for single-cell, single-mtDNA analysis, which allows simultaneous processing of hundreds of individual mtDNA molecules within >10,000 individual cells. Our microfluidic chip encapsulates individual cells in agarose beads, designed to have a sufficiently dense hydrogel network to retain mtDNA after lysis and provide a robust scaffold for subsequent multi-step processing and analysis. To mitigate the impact of the high viscosity of agarose required for mtDNA retention on the throughput of microfluidics, we developed a parallelized device, successfully achieving ~95% mtDNA retention from single cells within our microbeads at >700,000 drops/minute. To demonstrate utility, we analyzed specific regions of the single mtDNA using a multiplexed rolling circle amplification (RCA) assay. We demonstrated compatibility with both microscopy, for digital counting of individual RCA products, and flow cytometry for higher throughput analysis.

## Introduction

Mitochondria are sub-cellular organelles that play a key role in multiple cellular processes, such as bioenergetics, DNA repair, and control of cell cycle. ^[1]^ Each individual cell contains hundreds of mitochondria; the exact number varies depending on the cell type and the cell’s metabolic state. Each individual human mitochondrion has 1-10 copies of 16.6 kb double stranded circular DNA (mtDNA), which encodes 37 genes. ^[2]^ The mtDNA exhibits an estimated mutation rate approximately 5 to 15 times higher than the genomic DNA, ^[3]^ and large area deletions and single nucleotide variations (SNVs) in mtDNA have been discovered to be significant contributors to the onset and progression of a diverse set of conditions, including age-related diseases such as muscle weakness, neurodegenerative disorders, and cancer. ^[4]^ Key to understanding the role of these mutations is their distribution across the multiple mitochondria within each individual cell. For both healthy cells, and cells in a disease state, heteroplasmy has been observed, often involving regions within the non-coding mtDNA D-loop. ^[5]^ Roughly a quarter of the healthy population inherits a heteroplasmic mixture of both wild-type and variant mtDNA at birth, ^[6]^ and mtDNA mutations additionally accumulate with age. Within each individual cell, the number of mtDNA with mutations correlates with the cell’s pathological state. ^[7]^

To better understand mtDNA heteroplasmy there is a growing interest in developing new technologies to analyze the patterns of mtDNA mutation in individual mitochondrion within individual cells. This area of research offers a profound challenge to technologists, as it requires single cell, single organelle, and single molecule resolution. In particular, these assays require sufficient sensitivity to resolve mutations in single mtDNA and each single molecular measurement to be hierarchically associated with its cell of origin. Moreover, the platform must sample a sufficient number of mtDNA in each cell to detect rare heteroplasmic mutations, at a sufficient throughput to measure enough single cells to resolve the distribution of this heteroplasmy across diverse single cell populations. Conventional methodologies for detecting mutations in mtDNA lack the capability to effectively analyze mtDNA at the single-cell and single-mtDNA level. For instance, we recently adapted PCR-based approaches to achieve single-cell and single-mtDNA resolution using manual interventions such as microdissection, which restricted the number of cells and mitochondria that can be analyzed. ^[8]^ Others have used similar approaches to also analyze a limited number of mitochondria in a few cells. ^[9]^

Droplet microfluidic technologies have proven highly successful in single-cell analysis, ^[10]^ but have faced significant technological barriers for single-cell, single-organelle measurements ^[11]^ and thus have not been fully leveraged for single cell mtDNA heteroplasmy analysis. First, water-in-oil droplets do not easily allow for multi-step reactions with buffer exchanges including cell lysis, DNA purification, and signal amplification. ^[12]^ Secondly, water-in-oil emulsions are not compatible with conventional flow cytometers to read out the results of the assay with high-throughput, although recent work that use double emulsions has somewhat addressed this issue. ^[13]^ Agarose based droplet microfluidics have successfully addressed these issues for the analysis of genomic DNA. ^[14]^ Agarose is a naturally occurring hydrogel that can be reversibly solidified and melted, allowing agarose beads loaded with single cells to be generated using microfluidics at elevated temperatures and subsequently be solidified and transferred to an aqueous phase where multi-step assays can be carried out. The agarose forms a hydrogel network that can retain the macromolecules being studied while allowing reagents and cell lysate to transit into and out of the bead. Recently, rolling circle amplification (RCA) has been integrated with agarose microbeads for the multiplex digital measurement of nucleic acids. ^[15]^ These techniques allow amplification of single molecule signals within the hydrogel, allowing for digital counting of single molecules. ^[16]^ The size of RCA product is hundreds nm to a few µm, making it easily retained in the hydrogel network and thus compatible with these assays. ^[17]^ Despite these successes, these approaches have not yet been applied to the analysis of smaller macromolecules such as mtDNA because of the difficulty of retaining the ~100 nm sized mtDNA within a hydrogel network.

In this study, we have developed and validated an agarose-based droplet microfluidic methodology designed specifically for the analysis of single-cell, single-mitochondrial DNA (mtDNA) (**Fig. 1A**). Our approach allows hundreds of mtDNA molecules within over 10,000 individual cells to be individually analyzed, dramatically improving throughput compared to labor-intensive manual methods that have been applied to small cell populations (<100 cells) and only a few mtDNA per cell. ^[11]^ Our microchip encapsulates individual cells within 2% agarose beads, strategically formulated to have a sufficiently dense hydrogel network to ensure the retention of mtDNA during cellular lysis and multiple downstream processing and analysis steps (**Fig. 1B-D**). To address the challenge posed by the elevated viscosity of agarose necessary for optimal mtDNA retention and its impact on microfluidic throughput, a parallelized microfluidic chip was developed. This design achieved ~95% retention of mtDNA from single cells post-lysis within microbeads, operating at a throughput exceeding 700,000 drops per minute. The interrogation of distinct regions of the single mtDNA encapsulated within each bead was performed using fluorescence in situ hybridization (FISH) to label the RCA product that has been amplified from a padlock probe targeting a specific region of mtDNA (**Fig. 1E**). We demonstrated compatibility of this assay with both microscopy, enabling digital counting of individual RCA products, and flow cytometry for high-throughput analysis at the single cell level. Validation studies were performed using a multiplexed assay that targeted a common mtDNA large deletion region and total mtDNA content in individual cells (**Fig. 1F**). Additionally, we ensured minimal cross-contamination between beads by analyzing mtDNA from single cells from different species. This new agarose-based droplet microfluidic approach establishes a platform for high throughput single-cell, single-mtDNA mutation analysis, thereby opening new avenues for advancing our understanding of the biological role of mtDNA heteroplasmy.

**Figure 1.**
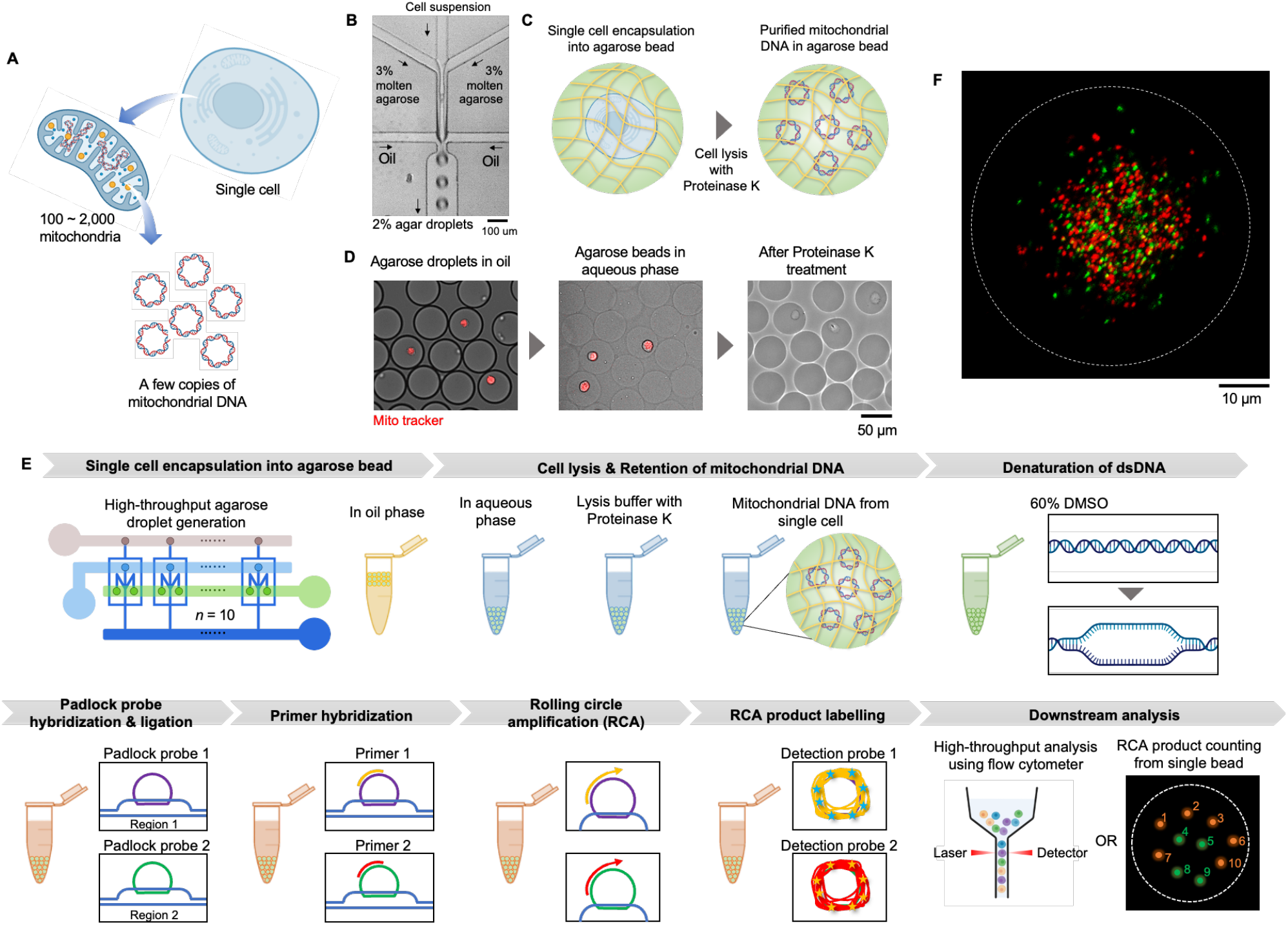
Schematic of high-throughput assay of single cell-single mtDNA. (A) Multiple mitochondria and mtDNA are present in a single cell, and some mitochondrial DNA exhibit mutations such as large area deletions. (B) Micrograph of microfluidic device for encapsulation of single cells into agarose beads. (C) After encapsulation of a single cell into an agarose bead, genomic material including mitochondrial DNA is retained in the agarose bead whereas protein is digested by proteinase K. (D) Cells are stained with MitoTracker and monitored after transferring to aqueous phase and Proteinase K treatment. (E) Sample preparation procedures for single cell-single mtDNA assay can be handled in a one-pot, high-throughput manner and can be analyzed by a confocal microscope or flow cytometer. (F) Fluorescence image showing the multiplex digital assay of mtDNA from single cell.

## Results

We experimentally evaluated the concentration of agarose necessary to retain mtDNA in agarose microbeads and found that 2% agarose was sufficient to retain ~100 nm (16.6 kb) sized mtDNA. ^[18]^ To this end, we first used isolated genomic material including mtDNA from cultured HEK293 cells (10^6^ cells) using a commercial genomic DNA isolation kit. We mixed the isolated genomic material (2.7 ng/µL) with various concentrations of molten agarose and used this aqueous phase to form emulsions (d = 100 µm) in fluorinated oil (2% Pico-surf in HFE-7500) using a flow focusing microfluidic droplet generator (**Figure S1A**). The advantage of microfluidics over vortexing based emulsification is improved precision and control over droplet diameter, which allows cells to be loaded into the droplets at precise concentrations to ensure a digital distribution of cells. To keep the agarose molten during the emulsification process, the temperature of molten agarose was kept at 50 °C using a combination of a space heater and syringe heaters (**Figure S2**). The agarose emulsion was collected in an ice bucket and subsequently incubated at 4 °C for 30 minutes to ensure gelation. After gelation of agarose droplets, the oil phase was removed followed by addition of 10% 1H,1H,2H,2H-perfluoro-1-octanol (PFO) in HFE-7500. Agarose beads were washed once with 1% Span80 in hexane followed by three washes with 0.1% Triton-X in TE buffer. Then, agarose beads were transferred to deionized water (DI) and kept in suspension for 24 hours (**Fig. 2A**), after which the mtDNA retained in the agarose was compared to that which had leaked into the DI using quantitative polymerase chain reaction (qPCR)(**Fig. 2B**). The relative quantity of mtDNA in agarose beads and supernatant was compared to assess mtDNA retention ratio. We found that as the agarose concentration was increased from 1-3% the quantity of mtDNA retained in the beads increased relative to that in the DI continuous phase. At 2% agarose the ratio reached 90.9%, which further improved marginally at 3% agarose (**Fig. 2C**).

**Figure 2.**
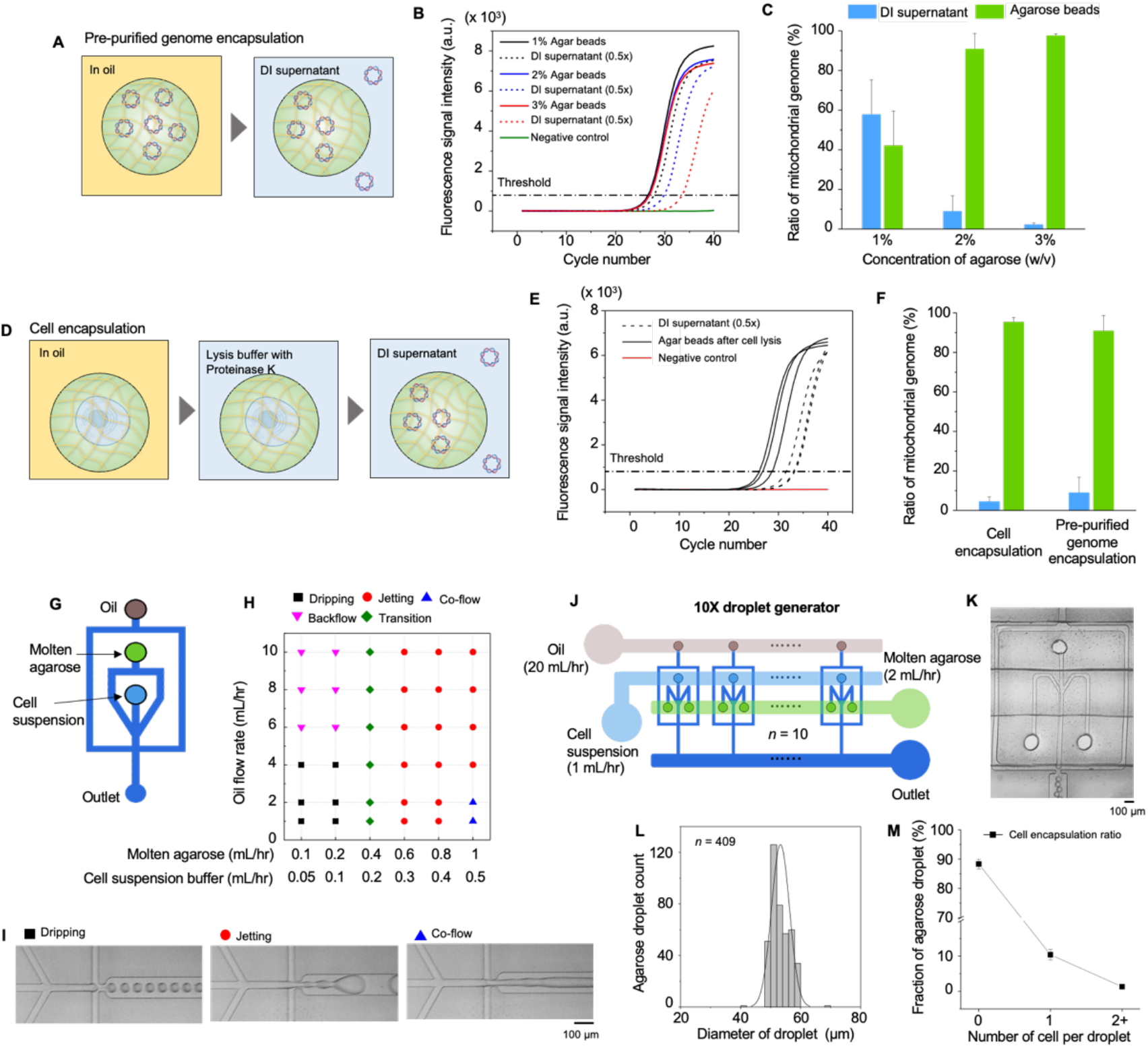
Validation for the retention of mtDNA in agarose beads. (A) A schematic showing the analysis method of pre-purified genome encapsulation experiment. (B) qPCR result for analyzing retention ratio of mtDNA depending on agarose concentration. (C) Retention ratio of mtDNA with respect to concentration of agarose beads (*n* = 3). (D) A schematic showing the analysis method of cell encapsulation experiment. (E) qPCR result of mitochondrial DNA retention in 2% agarose beads from cell encapsulation experiment. (F) Retention ratio of mtDNA was compared between cell and pre-purified genome encapsulation experiments (*n* = 3). (G) Schematic of the agarose droplet generator for single cell encapsulation. (H) Droplet generation regime transition plot depending on aqueous (molten agarose and. Cell suspension buffer) and oil phase flow rate. (I) Micrographs showing different droplet generation regimes. (J) Schematic of 10X parallelized high-throughput agarose droplet generator for single cell encapsulation. (K) Micrograph of a single unit of 10x agarose droplet generator. (L) Histogram of agarose droplet diameter generated by 10x droplet generator (*n* = 409). (M) Single cell encapsulation ratio into agarose droplet was plotted based on image analysis (*n* = 5 (total 2,177 agarose beads)). All error bars represent the standard deviation.

We next confirmed the concentration of agarose necessary to retain mtDNA from single cells loaded into the agarose beads and found similar results to isolated mtDNA. To this end, we loaded individual HEK293 cells into emulsions (d = 100 µm) of molten agarose suspended in fluorinated oil (2% Pico-surf in HFE-7500) using a flow focusing microfluidic droplet generator. To maintain cell viability until encapsulation in the agarose droplet that can be hindered by the high temperature (T > 50 °C) of molten agar, we employed a microfluidic droplet generator with separate inlets for molten agarose and cell suspension, wherein the cell suspension and agarose are mixed immediately prior to the droplet generation junction (**Figure S1B**). This design necessitated a high concentration of molten agarose (3%), to compensate for its dilution by the cell suspension, to achieve the same 2% agarose concentration in the microbeads as in the prior experiment. After encapsulation of single cell into 2% agarose droplet (0.2 mL/hr for 3% molten agar, 0.1 mL/hr for cell suspension, 2.5 mL/hr for fluorinated oil), the agarose droplet was solidified and transferred to lysis buffer (2 U/mL proteinase K and 0.5% lithium dodecyl sulfate (LDS) in TE buffer). After 30 minutes of incubation at 25 °C, lysis buffer was removed followed by washing with 2% T20 in DI, ethanol, and 0.02% T20 in DI. Then agarose beads were transferred to DI and kept in suspension for 24 hours (**Fig. 2D**), after which the mtDNA in the agarose beads was compared to that in the supernatant using qPCR (**Fig. 2E**). We achieved a ratio of mtDNA retained in the beads versus the supernatant outside the beads of 95.4% (**Fig. 2F**).

To isolate and retain mtDNA from single cells required the input of 3% agarose into our microfluidic chip, which has the unfortunate consequence of severely hindering microfluidic droplet generation throughput. The viscosity of agarose dramatically increases superlinearly as the concentration increases. ^[19]^ The 3% agarose has a viscosity of ~100 mPa⋅s (**Figure S3**), which causes the transition from dripping to jetting to occur at low flow rates (*ϕ*_d_ = 0.3 mL/hr for aqueous phase, *ϕ*_c_ = 2.5 mL/hr for the oil phase). It is in the dripping phase that the microfluidic droplet generator creates homogenous droplets (CV of diameter < 5%). Whereas in the dripping mode, droplets are highly inhomogenous (**Fig. 2G-I**). Due to the high viscosity of our dispersed phase, the maximum throughput that could be achieved in dripping mode on the single channel device was only 72,000 drops (51 µm, CV = 3%) per minute.

To increase throughput beyond the limit dictated by the viscous agar, we developed a parallelized microfluidic droplet generator fabricated using double side imprinting of PDMS ^[20]^. In prior work, as many as 1,000 generators have been parallelized by our group ^[20a]^, as well as similar work by others ^[21]^. To the same end, others have also employed a droplet splitting approach to enhance agarose droplet generation throughput that involves creating a large droplet and subsequently dividing it into eight smaller droplets. ^[22]^ In this study, we parallelized 10x generators (**Fig. 2J,K** and **Figure S4**), which provided sufficient throughput for this work. Notably, our design enables further parallelization that could increase throughput by another 100x, which is not possible in the droplet splitting approach. Our parallelized device generated agarose droplets with a diameter of 53 μm (CV = 5%) at a throughput of 720,000 drops/minute (**Fig. 2L**). We successfully loaded single cells into droplets. The ratio of agarose droplets with single cells was 10.4% while the ratio of empty droplets and droplets with two or more cells was respectively, 88.3% and 1.3% (N = 2,177), as predicted by Poisson statistics (**Fig. 2M**).

To perform single mtDNA analysis on single cells, we designed an RCA based assay that could be carried out within our agarose beads (**Fig. 3A**). RCA amplifies the signal from padlock probe annealed to a specific region within each mtDNA, by generating a long strand of DNA that can be subsequently labeled by multiple fluorescent labels using FISH. ^[23]^ Each RCA product is visualized as a punctum dispersed over the agarose bead which can be resolved using confocal fluorescence microscopy, ^[17a, 24]^ or quantified using flow cytometry with high-throughput. ^[25]^

**Figure 3.**
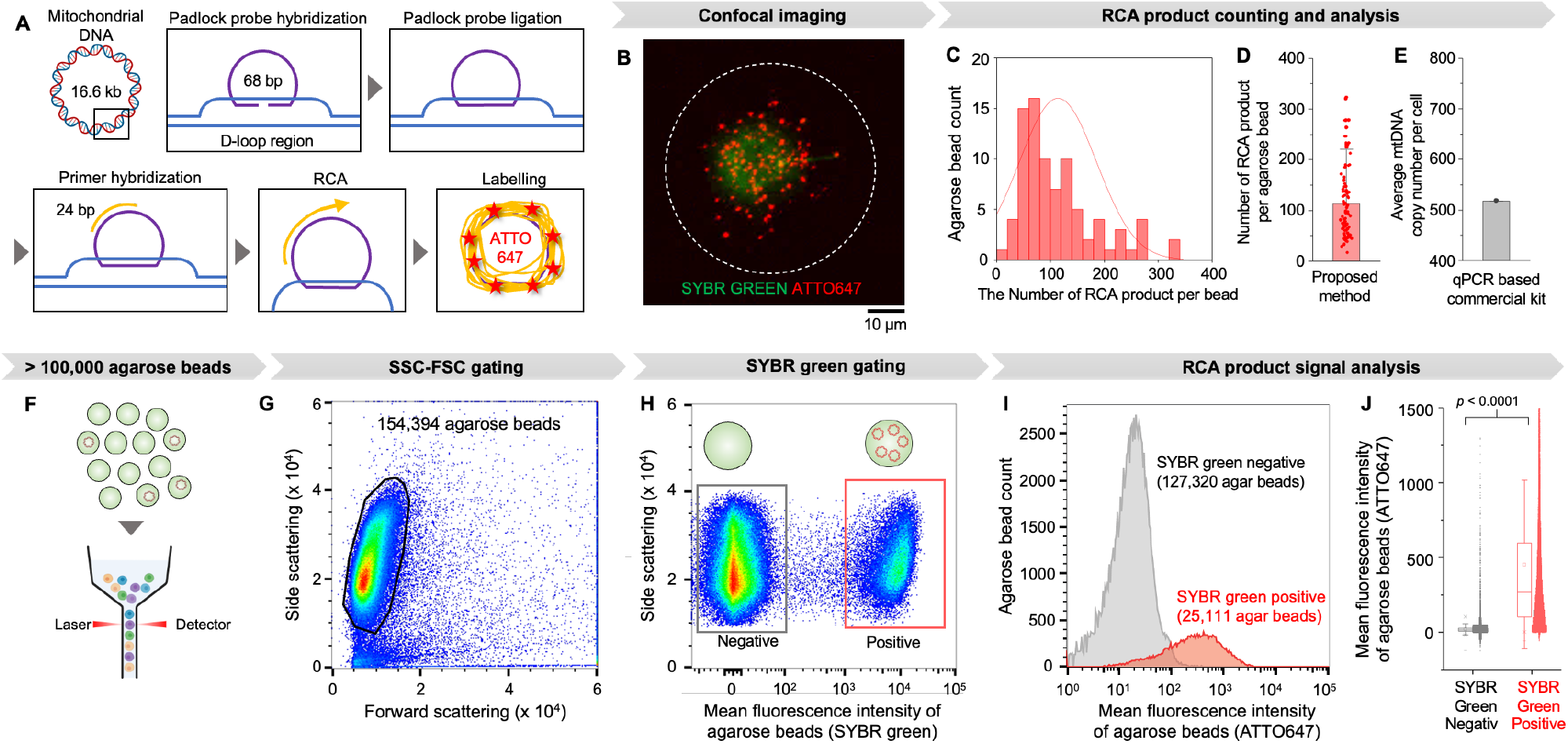
Single-cell, single-mtDNA assay using RCA. (A) Schematic showing the RCA based signal generation strategy from single mtDNA. (B) Image reconstructed from confocal microscope data showing the RCA product from single mtDNA (seen as red punctate) is distributed over the agarose bead whereas genomic DNA is concentrated at the center (seen as green center). (C) Histogram of single mitochondrial DNA counting from single cell (*n* = 83). (D) Distribution in the number of RCA products per agarose droplet (mitochondrial DNA copy number per single cell) shows high heterogeneity. An error bar represents the standard deviation. (E) Average mitochondrial DNA copy number per single cell was analyzed using a qPCR based commercialized kit. (F) High-throughput flow cytometry analysis of mtDNA from single cell. (G)154,394 agarose beads are gated from forward and side scattering gating. (H) Agarose beads with genomic material were gated by SYBR green stain as a positive control. (I) Positive beads on SYBR green gating showed brighter signal at ATTO647 emission channel that represents mtDNA copy number. (J) Mean fluorescence intensity distribution shows the heterogeneity of mitochondrial DNA copy number per single cell. The distributions were compared by one-way ANOVA. An error bar represents the standard deviation.

The workflow to perform our RCA based analysis is as follows, briefly. First, cells are loaded into ~50 µm agarose droplets using our parallelized microfluidic chip described above. The cells are loaded at a concentration such that on average there is <1 cell for every 10 droplets, following Poisson statistics, ensuring a digital distribution of cells in the droplets. The agarose droplets are then cooled to 4 ºC and transferred from oil to an aqueous solution. The cells are lysed within the agarose beads by suspending the beads in TE buffer with 2 U/mL proteinase K and 0.5% LDS for 30 minutes at 25 ºC. After lysis, the macromolecules, genomic material including the mtDNA, are retained. The beads are subsequently washed in 0.02% T20 in DI for 3 times. To target a mtDNA region of interest with padlock probes, the region needs to be single stranded. In a typical workflow, double stranded mtDNA is heated up to a high temperature (70 ºC) and rapidly cooled down to denature double stranded structures and maintain single strands. However, heat-based denaturation is not compatible with our agarose beads because agarose will melt at high temperatures, and mtDNA will be lost. To overcome this challenge, after cell lysis within the bead, the agarose beads are further incubated in 60% DMSO to denature double stranded structures. ^[26]^ After treating the agarose beads in 60% DMSO for 10 minutes and washing 3 times with 0.02% T20 in DI, the target region of denatured mtDNA is labelled with padlock probes and primers (**Figure S5**). First, padlock probes targeting the D-loop region of mtDNA are hybridized and ligated by T4 DNA ligase at 37 ºC for 1 hour. After washing unbound padlock probes, primers are hybridized to circular DNA annealed to the target region of mtDNA by incubation at 37 ºC for 30 minutes. After washing out unbound primers, RCA is performed at 37 °C for 2 hours along single-stranded circular padlock probes annealed to a specific region of mtDNA. Following RCA, the product is labeled with ATTO647 detection probes by washing out the RCA mixture and introducing the probes. We note that this multi-step amplification process, is readily carried out in our agarose beads using conventional laboratory equipment (i.e. centrifuge, pipettes), but would not be possible using water-in-oil droplet microfluidics, where robust, high throughput reagent exchange remains a technological challenge. ^[12]^

The RCA products from mtDNA in agarose beads are visualized and quantified using confocal microscopy. We found that mtDNA from the cell was distributed over the agarose bead, making it possible to visualize multiple signals from padlock probe annealed to mtDNA molecules (**Fig. 3B**). The number of RCA products in a single agarose bead was countable using confocal microscopy (Zeiss LSM 980). In addition to analyzing the mtDNA of each cell, we also incorporate SYBR green staining of the cellular genomic DNA (gDNA) to independently detect the beads that have indeed encapsulated a cell. gDNA (~3,000 Mb) is much larger than mtDNA (16.6 kb) and so is easily retained in the agarose bead. The RCA product on the D-loop region of mtDNA was stained with ATTO647 dye labeled FISH probes. Using the same HEK293 cells described above, on average, there were 113.8 RCA products per agarose bead (CV = 63%) (N = 83) (**Fig. 3C,D**). Compared to the result of qPCR-based mtDNA absolute copy number quantification kit (average 500 mtDNA per cell), fewer RCA products were counted using our method (**Fig. 3E**). There are two possibilities for the underestimated number of RCA products per bead. First, insufficient denaturation of the double-stranded mtDNA template could lead to incomplete strand separation, hindering efficient binding of the padlock probes, which would result in less mtDNA annealed with padlock probes. Second, the proximity of multiple mtDNA molecules within a single agarose bead may lead to aggregation of their respective RCA products (**Figure S6**), which can be further addressed by increasing mixing of mtDNA within microbeads before RCA or increasing the diameter of the microbeads.

While confocal microscopy has the advantage of digital counting of single RCA products within a single bead, the throughput of the analysis is low (1 agarose bead / 3 minutes). To increase the throughput of single cell, single mtDNA analysis, we demonstrate that our assay is compatible with flow cytometry. Because the agarose microbeads are <60 µm diameter and are suspended in an aqueous solution, mean fluorescence intensity (ATTO647) of each single agarose bead can be interrogated with conventional flow cytometer (Symphony A3 Lite (BD Biosciences) from Penn Cytomics & Cell Sorting Shared Resource Laboratory) without any further modifications to the assay described above (**Fig. 3F**). Based on forward and side scattering, single agarose beads were gated (**Fig. 3G**). Within that population, agarose beads that contain genomic material, based on SYBR green staining, and therefore had encapsulated a cell, were gated (**Fig. 3H**). Within this population, each bead’s fluorescence signal is quantified as a measure of one, or more, fluorescently labeled RCA products.

In our first demonstration, we analyzed the D-loop region in mtDNA in individual HEK293 human cells. The RCA product associated with the D-loop region of human mtDNA was labeled with ATTO647 dye. Compared to agarose beads that were not loaded with cells (negative in SYBR green signal labeling genomic DNA), agarose beads that did contain cells (positive in SYBR green signal) showed significantly greater fluorescence signal of ATTO647 labelled RCA product (*p* < 0.0001) (**Fig. 3I,J**). We analyzed the microbeads at a throughput of 11,280 agarose beads / minute, a throughput >10,000x faster than by imaging. Compatibility with both imaging and flow cytometry enables their combined application for high throughput sorting followed by imaging-based based downstream analysis. ^[27]^

We next evaluated whether mtDNA from one bead can transit between droplets using a duplex assay. To quantify potential cross-contamination we loaded our agarose beads with either CT26 mouse or K562 human cells. This assay verifies the specificity of our assay as well as minimal cross-contamination. A duplex assay was performed that labeled the D-loop regions of mouse and human mtDNA with ATTO565 and ATTO647, respectively (**Fig. 4A**). Without cross-contamination, each bead should contain either an ATTO565 or an ATTO647 signal, and the presence of beads positive for both these signals would indicate cross-contamination. First, the agarose beads were evaluated using fluorescence microscopy, and qualitatively revealed that there was no cross contamination between the microbeads (**Fig. 4B,C**). Cross-contamination was also quantified using flow cytometry, where we analyzed 3,165 cells (**Fig. 4D**). The beads containing human and mouse mtDNA were well separated, and could be clearly visualized by plotting the ratio (ATTO565 / ATTO647) of beads that contain cells (**Fig. 4E**). Moreover, we repeated the experiment using agarose beads containing only mouse or human mtDNA (**Fig. 4F,G**), to assess what the signal would look like with no possibility of cross-contamination, and these populations aligned with those identified in the above experiment described above.

**Figure 4.**
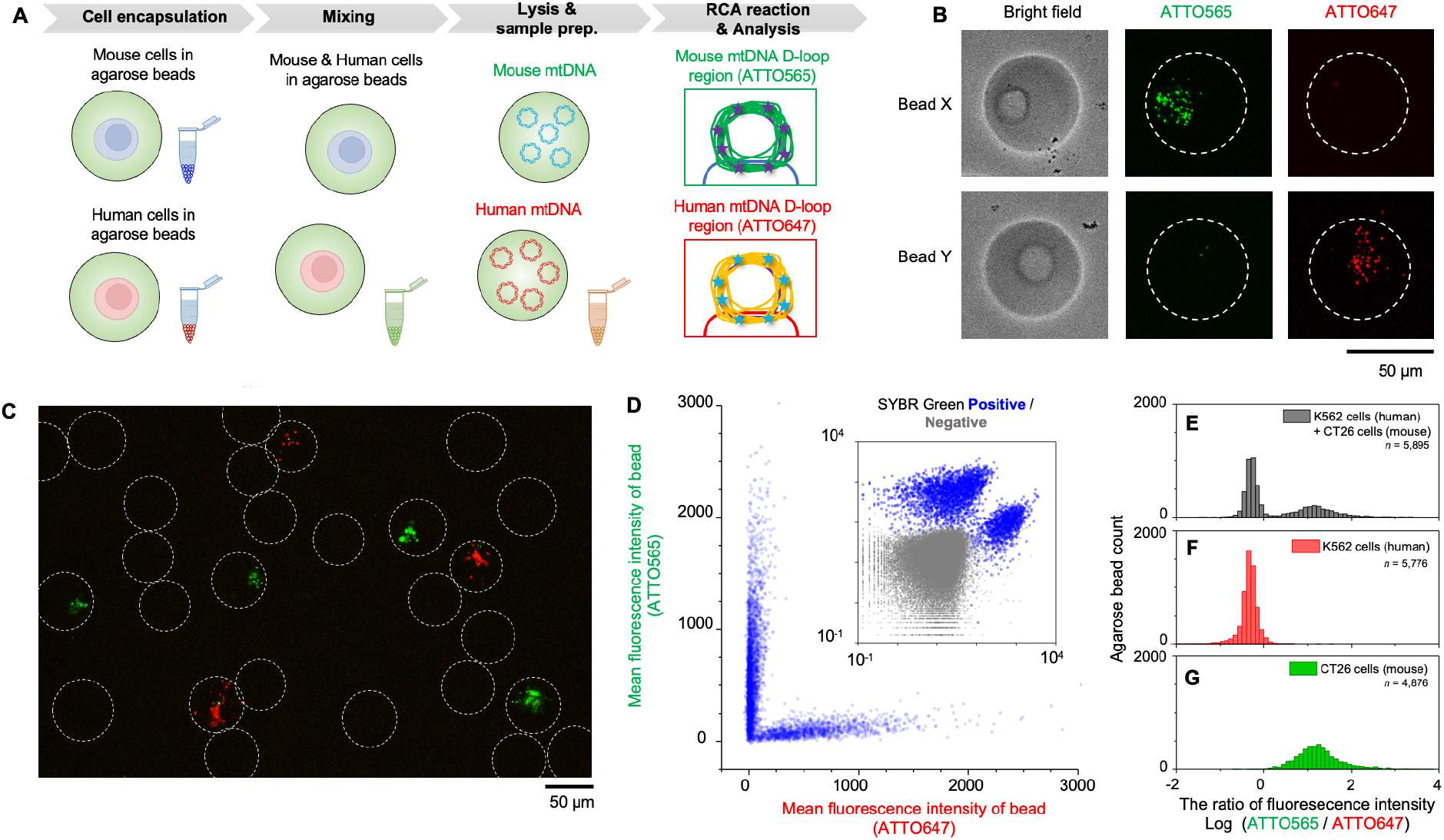
mtDNA cross-contamination assessment. (A) Schematic showing the experimental protocol using human and mouse cells. (B) Fluorescence micrographs of agarose beads with different fluorescence profile that show bead X containing a mouse cell and bead Y containing a human cell. (C) Large area fluorescence image of agarose beads that shows there is little overlap of fluorescence signal between agarose beads. (D) Scattering plot of the flow cytometry data showing mouse and human cells are distinguished by different fluorescence signal profiles (*n* = 3,165). (E) Histogram showing the ratio of fluorescence intensity (ATTO565 / ATTO647) of single agarose bead with genomic material. Control experiment with only (F) human or (G) mouse cells ensured each population originated from each species of cell.

We next demonstrated the capability of our assay to be multiplexed. Using multiple sets of padlock probes and primers, multiple regions of single mtDNA can be analyzed simultaneously at the single cell level. To evaluate this capability, we targeted two different regions in the mtDNA of single K562 human cells, including the 4,977 deletion area and the D-loop region (**Fig. 5A**). The 4,977 deletion is a common deletion that eliminates a region between 8,470 bp and 13,447 bp of the human mtDNA, and which is involved in various diseases such as myopathies, Alzheimer disease, and Kearns-Sayre syndrome, as it removes all 5 tRNA genes and 7 genes encoding Complex subunits. ^[28]^ Unlike 4,977 deletion region, D-loop region (positions 16,024 bp – 576 bp), the main non-coding area of the mtDNA and starting point of replication, does not show deletion. ^[29]^ Using fluorescence microscopy, we confirmed RCA products generated from the two different sets of padlock probes targeting D-loop and 4,977 deletion region within each bead (**Fig. 5B,C**). RCA products generated from padlock probes targeting the D-loop region were labeled with ATTO647 (red), while those from probes targeting the 4,977 deletion region were labeled with ATTO565 (green).

**Figure 5.**
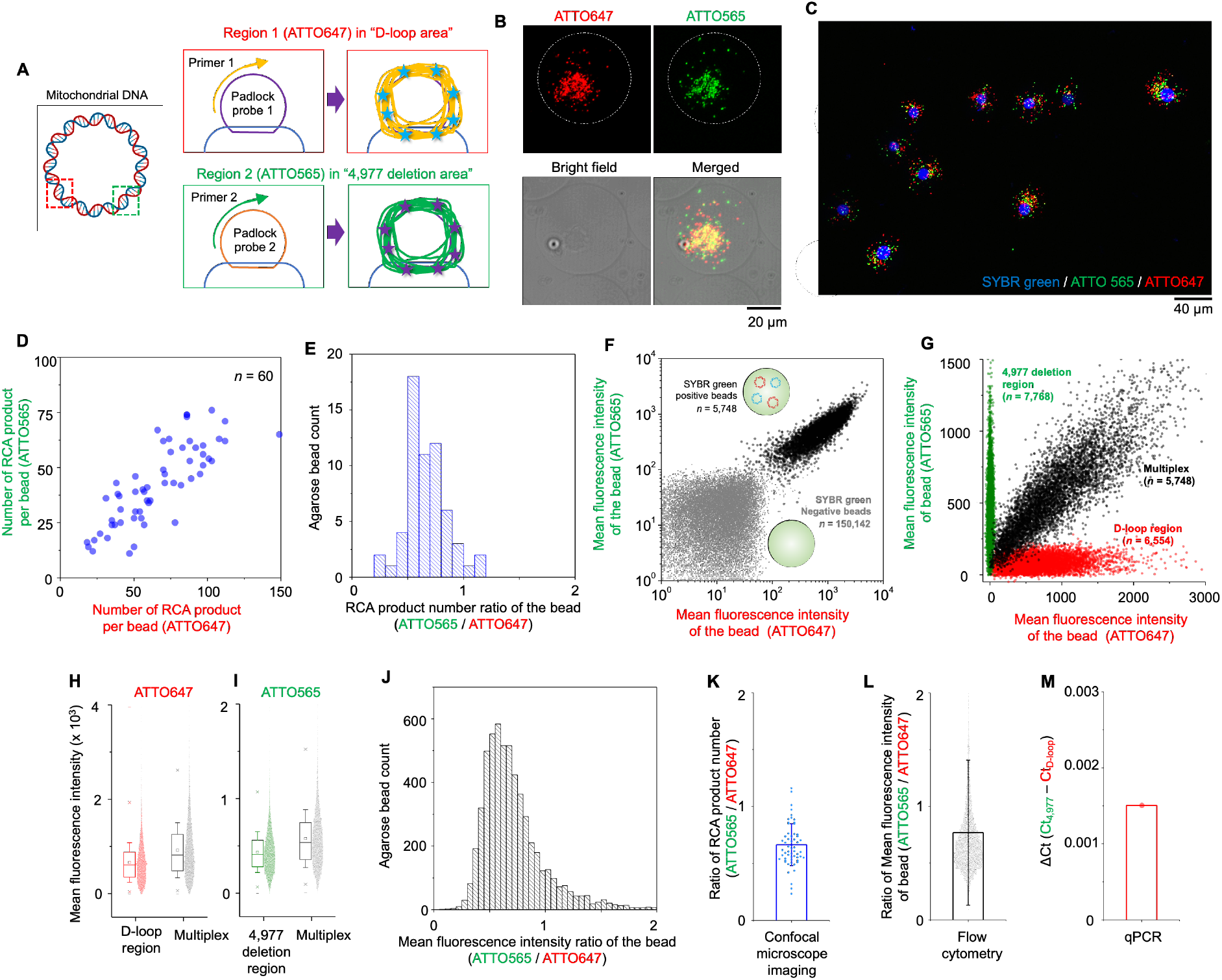
High-throughput multiplex analysis of single-cell mtDNA. (A) Schematic showing the multiplex analysis of single mtDNA where one RCA template targets conserved region and the other targets deletion region. (B) Fluorescence micrographs reconstructed from Z-projection of confocal microscope data showing the co-localization of both fluorescence signals in a single agarose bead. (C) Large area fluorescence image reconstructed from Z-projection of confocal microscope data. (D) Confocal imaging-based counting of two different RCA products from single agarose bead can be used for single-cell mtDNA deletion study (*n* = 60). (E) The ratio of the RCA product number labelled with ATTO565 or ATTO647 in a single agarose bead was plotted as a histogram. (F) Multiplex analysis of single-cell mtDNA was further quantified by flow cytometry where SYBR Green positive and negative beads are distinguished by both ATTO565 and ATTO647 fluorescence intensity. (G) Mean fluorescence intensity of agarose beads in multiplex analysis was compared to assay for each target region. (H) ATTO647 and (I) ATTO565 fluorescence intensity of assay for each target region was compared to the multiplex assay targeting both regions. Error bars represent the standard deviation. (J) High-throughput flow cytometer-based multiplex analysis of RCA products per bead for studying heteroplasmy of the mtDNA (*n* = 5,748). The ratio of fluorescence intensity of ATTO565 and ATTO647 from a single agarose bead was plotted. Plots showing the intracellular heterogeneity of mtDNA based upon (K) confocal imaging and (L) flow cytometry analysis. Error bars represent the standard deviation. (M) qPCR based bulk analysis of mtDNA 4,977 deletion that cannot resolve heterogeneity at a single-cell level.

We digitally quantified the RCA products generated by padlock probes targeting D-loop region and the 4,977 deletion area for individual K562 cells (N = 60) using imaging (**Fig. 5D**). Agarose beads that had encapsulated single cells were identified by positive SYBR Green gDNA labeling. On average, there were 66.65 (CV = 42%) RCA products labelled with ATTO647 (D-Loop) and 42.65 (CV = 42%) labeled with ATTO565 (4,977 deletion), each showing heterogeneity at the single cell level. To analyze heteroplasmy of mtDNA, the ratio of the number of RCA products at the 4,977 deletion area to that at the D-loop region can be quantified at the single cell level (**Fig. 5E**). The average ratio was 0.66 (CV = 28%). This ratio can be used as an indicator to screen cells with mtDNA 4,977 deletion (**Figure S7**).

To quantify the results of this multiplexed assay at high throughput, flow cytometry was incorporated to read out the multiplex digital assay of single cell-single mtDNA (**Fig. 5F**). As expected, we found that agarose beads with cells (SYBR Green +, N = 5,748) had significantly greater fluorescence intensity for RCA products associated with D-Loop and with the 4,977 deletion, compared to agarose beads without cells (SYBR Green -, N = 150,142). Additionally, we compared the signal from our multiplexed assay to assays targeting only 4,977 deletion region or only the D-loop region, and observed qualitative agreement (**Fig. 5G**). Moreover, we quantitatively compared the assay for D-loop region and 4,977 deletion region with the multiplex assay targeting both regions, to evaluate whether labeling with multiple probes had hindered the signal intensity of RCA products, and found no decrease in signal of multiplex assay (**Fig. 5H,I**). The variance measured using flow cytometry (CV= 63% for D-loop region and CV = 53% for 4,977 deletion region) (**Fig. 5F**) was similar to that measured using imaging (CV= 42% for D-loop region and CV = 42% for 4,977 deletion region) (**Fig 5E**). Variance in the flow cytometry measurement can be further reduced by eliminating all agarose beads containing more than one cell, by decreasing the ratio of cells to beads, and eliminating ruptured cells that can have fewer mtDNA.

Heteroplasmy within individual cells was analyzed in flow cytometry by quantifying the fluorescence intensity ratio of ATTO565 (4,977 deletion region) and ATTO647 (D-loop region) was analyzed for 5,748 individual cells (**Fig. 5J**). The average ratio was 0.77 (CV = 83%). This ratio can be used to identify cells that have a particular level of heteroplasmy for downstream analysis. ^[27]^ The ratio of RCA products number per bead or mean fluorescence signal intensity ratio of a bead are analyzable with imaging method (**Fig. 5K**) or flow cytometry based method (**Fig. 5L**), which successfully resolved heteroplasmy that cannot be resolved using gold standard qPCR based bulk analysis (**Fig. 5M**).

## Discussion

We have developed and validated a high-throughput method for single cell-single mtDNA digital assay based on microfluidic-generated agarose microbeads. Our approach achieved >95% of mtDNA retention ratio in agarose beads, and allowed an RCA-based multistep analysis of >10,000 single cells processed in a single tube with minimal cross-contamination. A high-throughput, parallelized agarose droplet generator was developed that achieved a throughput more than sufficient for this study (> 700,000 droplets / min), and was designed to be readily scaled to orders of magnitudes greater throughputs to analyze even larger numbers of cells. RCA products from single cell-mtDNA were analyzed using both confocal microscope imaging and flow cytometry. Using flow cytometry, >2,500,000 mtDNA of >25,000 cells were analyzed with a throughput of 1,200 cells per minute. Single-cell mtDNA copy number assay was validated by analyzing D-loop region of mtDNA, and given its known role as a biomarker of various diseases linked to cellular energy production, the measured mtDNA copy number per cell holds promise as a potential indicator for disease diagnosis and monitoring by overcoming heterogeneity. ^[30]^ The multiplex assay was further validated by analyzing both D-loop and 4,977 deletion region of mtDNA, which confirmed the heterogeneity of mtDNA across single cells and heteroplasmy within a single cell. While prior droplet microfluidics-based approaches to single-cell analysis of transposase-accessible chromatin with sequencing (scATAC-seq) have been used to infer mtDNA heteroplasmy computationally, ^[31]^ our method is able to directly measure single cell heterogeneity and mtDNA heteroplasmy by quantifying copy number of mtDNA from a single cell, by counting RCA products using microscopy or quantifying fluorescence signal using flow cytometry with high-throughput. Another advantage of our method is its applicability to all mtDNA regions, including those beyond the reach of transposase-accessible chromatin sequencing, which opens exciting possibilities for studying heteroplasmy arising from mtDNA deletions as well as mitochondrial SNVs.

One aspect of our work, for which there is a clear opportunity for future improvement is the under-estimation of mtDNA copy number compared to qPCR based conventional bulk analysis method. The average number of mtDNA measured using our RCA approach was approximately only 20% of the average mtDNA copy number per cell detected by a qPCR based commercial kit (**Fig. 3D,E**). We reasoned that this difference was unlikely due to leakage of the mtDNA from the beads, because we had independently quantified that retention rate to be 95%. Therefore, we suspect that it is the efficiency of labeling the mtDNA with padlock probes that is leading to this underestimation. We used DMSO to denature mtDNA, which can renature upon subsequent washing and buffer exchange procedures including salts. ^[26]^ In future work, we can instead use temperature based methods for denaturing the double stranded DNA, which have higher efficiency, by using heat-resistant hydrogels, such as polyacrylamide or alginate. ^[32]^

Our method for high-throughput single cell, single mitochondrial DNA assay using hydrogel droplet microfluidics has potential as a tool for both mechanistically studying mtDNA mutation-associated diseases for understanding disease threshold of deleterious SNVs and deletions through comparison with healthy control samples, as well as a diagnostic for screening patient cells for mtDNA changes. Furthermore, as our method is compatible with flow cytometry, higher dimensional analysis at the single cell level is possible. Agarose beads that contain single cell genomic material (gDNA and mtDNA) can be measured and sorted based on predefined criteria for further analysis by sequencing or imaging methods. There is opportunity to greatly expand the multiplexing capability of our technology beyond the 2x demonstrated in this work to many more regions of the mtDNA. Firstly, additional dyes can be used in FISH labeling of additional distinct RCA products. Beyond that, multiplexing can be further increased by labeling each RCA products with distinct combinations of dyes, which can be readout using microscopy. ^[33]^ Additionally, the method can be applied to analyze single cells isolated from tissue biopsies, enabling studies on the role of mtDNA mutations in various diseases. ^[34]^ As the principle of single cell-mtDNA retention on agarose beads is based on size exclusion, it can be applied to the analysis of smaller sized genomic material belonging to other subcellular organelles or viruses by engineering the pore structure of hydrogel or utilizing a hydrogel with a smaller pore size. ^[35]^

## Supporting information

Methods, Figure S1–S7, and Table S1–2 are available in supporting information.

## Acknowledgements

The authors would like to acknowledge Sabine Baxter for her assistance to culture mammalian cells and Hanfei Shen for his helpful discussion and feedback. This work is supported by the following funding sources: National Human Genome Research Institute (RM1-HG-010023), National Cancer Institute (R21CA236653, R33CA278551), National Institute of Mental Health (R33-NIMH-118170), and National Institute of Allergy and Infectious Diseases (R33-AI-147406)

## References

[1] a) D. D. Newmeyer, S. Ferguson-Miller, Cell 2003, 112, 481–490; b) J. R. Friedman, J. Nunnari, Nature 2014, 505, 335–343.

[2] P. A. Gammage, C. Frezza, BMC Biology 2019, 17, 53.

[3] B. A. I. Payne, I. J. Wilson, P. Yu-Wai-Man, J. Coxhead, D. Deehan, R. Horvath, R. W. Taylor, D. C. Samuels, M. Santibanez-Koref, P. F. Chinnery, Human Molecular Genetics 2012, 22, 384–390.

[4] a) T. Chen, J. He, Y. Huang, W. Zhao, Journal of Human Genetics 2011, 56, 689–694; b) S. Vyas, E. Zaganjor, M. C. Haigis, Cell 2016, 166, 555–566; c) K. Zhang, R. Deng, X. Teng, Y. Li, Y. Sun, X. Ren, J. Li, Journal of the American Chemical Society 2018, 140, 11293–11301.

[5] J. B. Stewart, P. F. Chinnery, Nature Reviews Genetics 2015, 16, 530–542.

[6] S. Luo, C. A. Valencia, J. Zhang, N.-C. Lee, J. Slone, B. Gui, X. Wang, Z. Li, S. Dell, J. Brown, S. M. Chen, Y.-H. Chien, W.-L. Hwu, P.-C. Fan, L.-J. Wong, P. S. Atwal, T. Huang, Proceedings of the National Academy of Sciences 2018, 115, 13039–13044.

[7] a) K. Ye, J. Lu, F. Ma, A. Keinan, Z. Gu, Proceedings of the National Academy of Sciences 2014, 111, 10654–10659; b) Y. Wang, X. Guo, K. Ye, M. Orth, Z. Gu, Proceedings of the National Academy of Sciences 2021, 118, e2014610118.

[8] J. Morris, Y.-J. Na, H. Zhu, J.-H. Lee, H. Giang, A. V. Ulyanova, G. H. Baltuch, S. Brem, H. I. Chen, D. K. Kung, T. H. Lucas, D. M. O’Rourke, J. A. Wolf, M. S. Grady, J.-Y. Sul, J. Kim, J. Eberwine, Cell Reports 2017, 21, 2706–2713.

[9] a) L. He, P. F. Chinnery, S. E. Durham, E. L. Blakely, T. M. Wardell, G. M. Borthwick, R. W. Taylor, D. M. Turnbull, Nucleic Acids Research 2002, 30, e68–e68; b) N. R. Phillips, M. L. Sprouse, R. K. Roby, Scientific Reports 2014, 4, 3887; c) K. A. Rygiel, J. P. Grady, R. W. Taylor, H. A. L. Tuppen, D. M. Turnbull, Scientific Reports 2015, 5, 9906; d) R. O’Hara, E. Tedone, A. Ludlow, E. Huang, B. Arosio, D. Mari, J. W. Shay, Genome Research 2019, 29, 1878–1888; e) C. Bi, L. Wang, Y. Fan, B. Yuan, G. Ramos-Mandujano, Y. Zhang, S. Alsolami, X. Zhou, J. Wang, Y. Shao, P. Reddy, P.-Y. Zhang, Y. Huang, Y. Yu, Juan C. Izpisua Belmonte, M. Li, Nucleic Acids Research 2023, 51, e48–e48.

[10] a) D.-K. Kang, M. Monsur Ali, K. Zhang, E. J. Pone, W. Zhao, TrAC Trends in Analytical Chemistry 2014, 58, 145–153; b) B. Li, X. Ma, J. Cheng, T. Tian, J. Guo, Y. Wang, L. Pang, Frontiers in Bioengineering and Biotechnology 2023, 11.

[11] M. J. Siedlik, Z. Yang, P. S. Kadam, J. Eberwine, D. Issadore, Small 2021, 17, 2005793.

[12] M. J. Siedlik, D. Issadore, Microsystems & Nanoengineering 2022, 8, 46.

[13] a) K. K. Brower, C. Carswell-Crumpton, S. Klemm, B. Cruz, G. Kim, S. G. K. Calhoun, L. Nichols, P. M. Fordyce, Lab on a Chip 2020, 20, 2062–2074; b) M. Khariton, C. J. McClune, K. K. Brower, S. Klemm, E. S. Sattely, P. M. Fordyce, B. Wang, Analytical Chemistry 2023, 95, 935–945.

[14] a) R. Novak, Y. Zeng, J. Shuga, G. Venugopalan, D. A. Fletcher, M. T. Smith, R. A. Mathies, Angewandte Chemie International Edition 2011, 50, 390–395; b) F. Lan, B. Demaree, N. Ahmed, A. R. Abate, Nature Biotechnology 2017, 35, 640–646; c) Y. Nishikawa, M. Kogawa, M. Hosokawa, R. Wagatsuma, K. Mineta, K. Takahashi, K. Ide, K. Yura, H. Behzad, T. Gojobori, H. Takeyama, ISME Communications 2022, 2, 92.

[15] K. Sato, Y. Kitamura, N. Sasaki, K. Mawatari, M. Nilsson, T. Kitamori, Proceeding of 14th International Conference on Miniaturized Systems for Chemistry and Life Sciences, 3 - 7 October 2010, Groningen, The Netherlands, 2010.

[16] a) S. C. Chapin, P. S. Doyle, Analytical Chemistry 2011, 83, 7179–7185; b) A. Rakszewska, R. J. Stolper, A. B. Kolasa, A. Piruska, W. T. S. Huck, Angewandte Chemie International Edition 2016, 55, 6698–6701.

[17] a) J. Björkesten, S. Patil, C. Fredolini, P. Lönn, U. Landegren, Nucleic Acids Research 2020, 48, e73–e73; b) R. R. G. Soares, N. Madaboosi, M. Nilsson, Accounts of Chemical Research 2021, 54, 3979–3990.

[18] C. Kukat, C. A. Wurm, H. Spåhr, M. Falkenberg, N.-G. Larsson, S. Jakobs, Proceedings of the National Academy of Sciences 2011, 108, 13534–13539.

[19] M. Ghebremedhin, S. Seiffert, T. A. Vilgis, Current Research in Food Science 2021, 4, 436–448.

[20] a) H.-H. Jeong, V. R. Yelleswarapu, S. Yadavali, D. Issadore, D. Lee, Lab on a Chip 2015, 15, 4387–4392; b) Z. Yang, Y. Atiyas, H. Shen, M. J. Siedlik, J. Wu, K. Beard, G. Fonar, J. P. Dolle, D. H. Smith, J. H. Eberwine, D. F. Meaney, D. A. Issadore, Nano Letters 2022, 22, 4315–4324.

[21] a) T. Nisisako, T. Ando, T. Hatsuzawa, Lab on a Chip 2012, 12, 3426–3435; b) M. B. Romanowsky, A. R. Abate, A. Rotem, C. Holtze, D. A. Weitz, Lab on a Chip 2012, 12, 802–807.

[22] L. Liu, C. K. Dalal, B. M. Heineike, A. R. Abate, Lab on a Chip 2019, 19, 1838–1849.

[23] a) M. M. Ali, F. Li, Z. Zhang, K. Zhang, D.-K. Kang, J. A. Ankrum, X. C. Le, W. Zhao, Chemical Society Reviews 2014, 43, 3324–3341; b) P. S. Kadam, Z. Yang, Y. Lu, H. Zhu, Y. Atiyas, N. Shah, S. Fisher, E. Nordgren, J. Kim, D. Issadore, J. Eberwine, Submitted Manuscript.

[24] J. Zhang, J. Shi, H. Zhang, Y. Zhu, W. Liu, K. Zhang, Z. Zhang, Journal of Extracellular Vesicles 2020, 10, e12025.

[25] a) Z. Hu, F. Xu, G. Sun, S. Zhang, X. Zhang, Chemical Communications 2020, 56, 5409–5412; b) C. Wu, T. J. Dougan, D. R. Walt, ACS Nano 2022, 16, 1025–1035.

[26] X. Wang, H. J. Lim, A. Son, Environ Health Toxicol 2014, 29, e2014007.

[27] J. de Rutte, R. Dimatteo, M. M. Archang, M. van Zee, D. Koo, S. Lee, A. C. Sharrow, P. J. Krohl, M. Mellody, S. Zhu, J. V. Eichenbaum, M. Kizerwetter, S. Udani, K. Ha, R. C. Willson, A. L. Bertozzi, J. B. Spangler, R. Damoiseaux, D. Di Carlo, ACS Nano 2022, 16, 7242–7257.

[28] A. A. M. Yusoff, W. S. W. Abdullah, S. Khair, S. M. A. Radzak, Oncol Rev 2019, 13, 409.

[29] a) H. Sharma, A. Singh, C. Sharma, S. K. Jain, N. Singh, Cancer Cell International 2005, 5, 34; b) T. J. Nicholls, M. Minczuk, Experimental Gerontology 2014, 56, 175–181.

[30] C. A. Castellani, R. J. Longchamps, J. Sun, E. Guallar, D. E. Arking, Mitochondrion 2020, 53, 214–223.

[31] C. A. Lareau, L. S. Ludwig, C. Muus, S. H. Gohil, T. Zhao, Z. Chiang, K. Pelka, J. M. Verboon, W. Luo, E. Christian, D. Rosebrock, G. Getz, G. M. Boland, F. Chen, J. D. Buenrostro, N. Hacohen, C. J. Wu, M. J. Aryee, A. Regev, V. G. Sankaran, Nature Biotechnology 2021, 39, 451–461.

[32] M. V. Tamminen, M. P. J. Virta, Frontiers in Microbiology 2015, 6.

[33] M. J. Levesque, A. Raj, Nature Methods 2013, 10, 246–248.

[34] a) Á. Quintanal-Villalonga, J. M. Chan, I. Masilionis, V. R. Gao, Y. Xie, V. Allaj, A. Chow, J. T. Poirier, D. Pe’er, C. M. Rudin, L. Mazutis, STAR Protocols 2022, 3, 101776; b) D. Soteriou, M. Kubánková, C. Schweitzer, R. López-Posadas, R. Pradhan, O.-M. Thoma, A.-H. Györfi, A.-E. Matei, M. Waldner, J. H. W. Distler, S. Scheuermann, J. Langejürgen, M. Eckstein, R. Schneider-Stock, R. Atreya, M. F. Neurath, A. Hartmann, J. Guck, Nature Biomedical Engineering 2023, 7, 1392–1403.

[35] a) Tissue Engineering Part B: Reviews 2010, 16, 371–383; b) R. Foudazi, R. Zowada, I. Manas-Zloczower, D. L. Feke, Langmuir 2023, 39, 2092–2111.

